# Neural mechanisms underlying distractor inhibition on the basis of feature and/or spatial expectations

**DOI:** 10.1101/2020.04.05.026070

**Authors:** Dirk van Moorselaar, Nasim Daneshtalab, Heleen A. Slagter

**Author notes:** Please address correspondence to: Dr. Dirk van Moorselaar, Department of Experimental and Applied Psychology, Vrije Universiteit Amsterdam.

## Abstract

A rapidly growing body of research indicates that inhibition of distracting information may not be under flexible, top-down control, but instead heavily relies on expectations derived from past experience about the likelihood of events. Yet, how expectations about distracting information influence distractor inhibition at the neural level remains unclear. To determine how expectations induced by distractor features and/or location regularities modulate distractor processing, we measured EEG while participants performed two variants of the additional singleton paradigm. Critically, in these different variants, target and distractor features either randomly swapped across trials, or were fixed, allowing for the development of distractor feature-based expectations. Moreover, the task was initially performed without any spatial regularities, after which a high probability distractor location was introduced. Our results show that both distractor feature- and location regularities contributed to distractor inhibition, as indicated by corresponding reductions in distractor costs during visual search and an earlier distractor-evoked Pd component. Yet, control analyses showed that while observers were sensitive to regularities across longer time scales, the observed effects to a large extent reflected intertrial repetition. Large individual differences further suggest a functional dissociation between early and late Pd components, with the former reflecting early sensory suppression related to intertrial priming and the latter reflecting suppression sensitive to expectations derived over a longer time scale. Also, counter to some previous findings, no increase in anticipatory alpha-band activity was observed over visual regions representing the expected distractor location, although this effect should be interpreted with caution as the effect of spatial statistical learning was also less pronounced than in other studies. Together, these findings suggest that intertrial priming and statistical learning may both contribute to distractor suppression and reveal the underlying neural mechanisms.

## 1. Introduction

Our visual environment is filled with distractions, ranging from billboards alongside the road to the blinking light on your smart phone demanding your attention. Whether such visual distractors automatically capture attention in a stimulus-driven fashion (Theeuwes, 2010), or alternatively, only capture attention in a goal-driven manner when they match the current attentional set (e.g., (Folk, Remington, & Johnston, 1992), has long been a topic of active debate. Yet, growing evidence highlights that this theoretical dichotomy cannot account for many situations in which strong attentional biases arise from previous selection episodes (Awh, Belopolsky, & Theeuwes, 2012). As a specific case, there are now many demonstrations that location-probability learning of not only the target but also of the distractor benefits visual search, such that salient distractors are more efficiently ignored at locations with a high distractor probability (Ferrante et al., 2018; Goschy, Bakos, Müller, & Zehetleitner, 2014; Leber, Gwinn, Hong, & O’Toole, 2016; Wang & Theeuwes, 2018b). Critically, such distractor suppression is not observed when the upcoming distractor location is cued on a trial-by-trial basis (Noonan et al., 2016; Wang & Theeuwes, 2018a). Together, these findings suggest that distractor inhibition may not be under voluntary, top-down control, but instead greatly relies on statistical learning or expectations derived from past experience about the certainty or likelihood of events (Noonan, Crittenden, Jensen, & Stokes, 2018; Theeuwes, 2019; van Moorselaar & Slagter, 2020).

Although there is now a general consensus that learning, either implicitly or explicitly, about the spatial distribution of salient, but task-irrelevant distractors in a search display minimizes their interference, the processing stage at which this learned suppression is realized appears flexible. Whereas in some visual search studies the distractor-location effect (i.e., reduced distractor interference at high- vs. low-probability distractor location) was accompanied by impaired target processing at high probability distractor locations, in other studies target processing was unaffected by the spatial distractor regularity across searches. A notable difference between these studies is that in many of the former studies, the defining target and distractor features swapped randomly across trials (e.g., (Ferrante et al., 2018; Wang & Theeuwes, 2018b), such that learning could necessarily only occur on the spatial level, whereas in the latter studies, the color of the distractor was fixed across trials, hence also allowing for learning at the feature level (Allenmark, Zhang, Liesefeld, Shi, & Müller, 2019; Sauter, Liesefeld, Zehetleitner, & Müller, 2018; Zhang, Allenmark, Liesefeld, Shi, & Muller, 2019). This dissociation suggests that spatially specific suppression can shift from generic to feature-specific, when regularities across displays allow for learning at the both the spatial and the feature level (Failing, Feldmann-Wustefeld, Wang, Christian, & Theeuwes, 2019)(but see (Heinrich R Liesefeld & Müller, 2020). Here, we set out to determine how spatial and feature expectations independently and/or in interaction influence distractor processing and interference at the neural level.

Previous event-related potential (ERP) studies support the notion that distractor learning at either the feature or the spatial level can prevent attentional capture by salient distractors. In search displays with randomly mixed distractor (vis-à-vis non-distractor) features (i.e., shape and color), salient distractors typically elicit an N2pc (Burra & Kerzel, 2013), an ERP index of attentional selection (Luck & Hillyard, 1994b), especially so after feature swaps (Hickey, Olivers, Meeter, & Theeuwes, 2011)(but see)(McDonald, Green, Jannati, & Di Lollo, 2013). By contrast, in fixed-feature variants of the same tasks, which allow for the development of feature expectations, the same distractors elicit a Pd, an ERP index linked to inhibition (Gaspelin & Luck, 2018b; Sawaki & Luck, 2010), indicating that salient distractors were effectively filtered out before they could capture attention (Burra & Kerzel, 2013; Jannati, Gaspar, & McDonald, 2013). Similarly, Wang, van Driel, Ort, and Theeuwes (2019) showed that at high probability distractor locations, singleton distractors elicited a Pd, whereas without this spatial bias (measured in an independent set of participants), the same distractors first elicited an N2pc. Thus, expectations at either the feature or spatial level are by itself sufficient to implement pre-attentive distractor suppression, raising the question how distractor suppression develops and is neurally implemented when feature and spatial expectations can be combined.

As outlined above, behavioral evidence suggests that generic spatial suppression only arises when the spatial distractor regularity cannot be combined with feature expectations. Consistent with this, in the Wang et al. (2019) study not only distractors, but also targets elicited a Pd at the high probability distractor location. What is more, both distractors and targets at high probability distractor locations also elicited an early Pd component, indicative of early suppression (Weaver, van Zoest, & Hickey, 2017). These findings suggest that learning-dependent spatial suppression was set up in advance in a feature blind manner. Consistent with this, alpha-band activity, a marker of top-down neural inhibition (Jensen & Mazaheri, 2010), tuned to the high probability distractor location already increased before search display onset. While such anticipatory and generic spatial suppression may be the relatively most optimal strategy under conditions in which only information about the likely distractor location is available, it is obviously more advantageous from an ecological perspective to selectively suppress only distractors based on their specific properties. In this respect it is notable that two recent EEG studies did not observe any changes in anticipatory alpha-band activity as a function of distractor location learning when target and distractor features did not swap across trial (Noonan et al., 2016; van Moorselaar & Slagter, 2019). Here, we therefore also examined anticipatory alpha-band modulations and target processing at high probability distractor locations in conditions where spatial expectations could or could not be combined with feature expectations, in addition to directly contrasting condition effects on distractor processing using ERPs.

Specifically, we measured EEG while observers performed, in separate sessions, fixed-feature and mixed-feature variants of the additional singleton paradigm (Theeuwes, 1992). In the fixed-feature version, distractor and target features were fixed throughout an experimental block and participants could therefore develop expectations at the feature level. By contrast, in the mixed-feature version, the shapes and colors of the target and distractors randomly swapped across trials so that participants could not form any predictions at the feature level (see Figure 1). Critically, in both variants, halfway through the experiment, and unbeknownst to the participant, we introduced a high probability distractor location. This allowed us to examine how feature and spatial expectations may interact in modulating distractor processing and interference. Without any spatial regularities we expected distractors to elicit a Pd in the fixed-feature variant, but an N2pc, possibly followed by a subsequent Pd signaling reactive distractor inhibition (Heinrich René Liesefeld, Liesefeld, Töllner, & Müller, 2017; van Moorselaar & Slagter, 2019), in the mixed-feature variant. By contrast, with a spatial distractor bias we expected a distractor Pd irrespective of the feature variant, and no distractor-evoked N2pc. Yet, whether this Pd differed between feature variants was an open question. Moreover, following Wang et al. (2019), we expected a feature-blind Pd (i.e., a Pd elicited by both targets and distractors) at high probability distractor locations and increased anticipatory alpha power contralateral to the high probability distractor location, both indicative of generic spatial suppression, in the mixed-feature variant, but not in the fixed-feature variant, when expectations about distractor features could also be formed.

**Figure 1.**
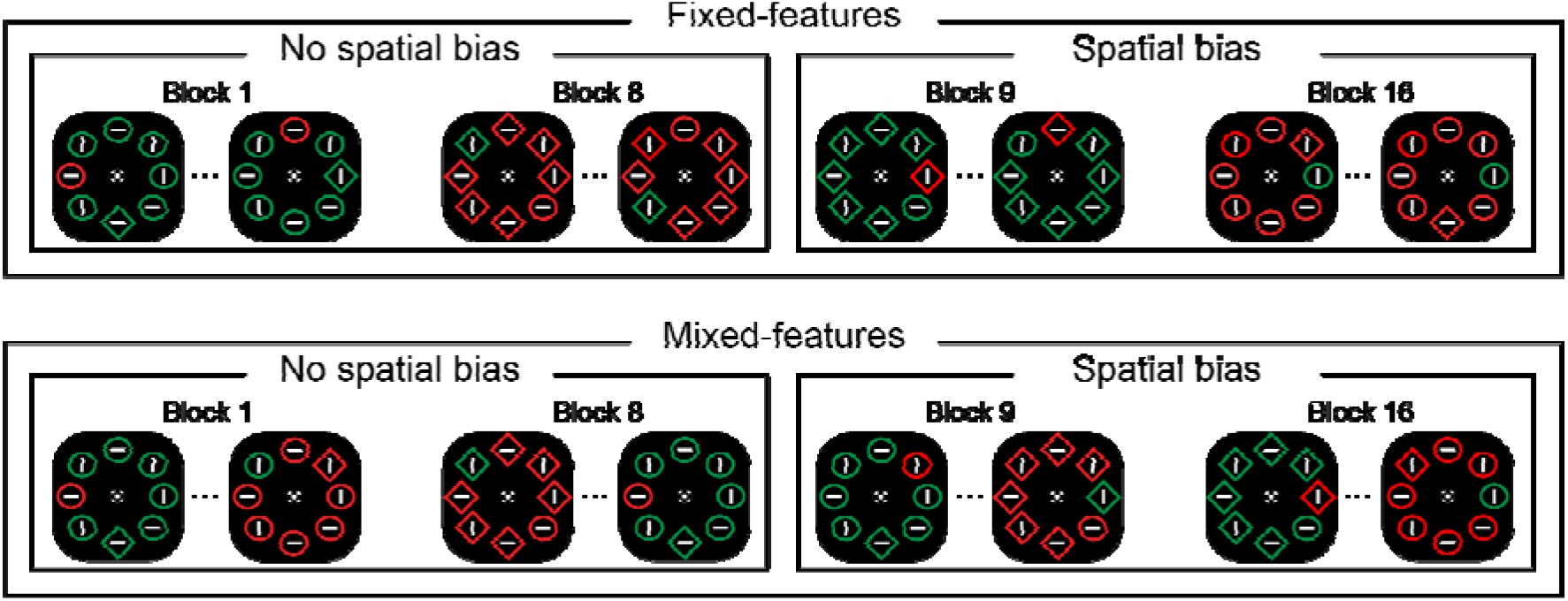
Experimental paradigm. In each trial participants had to indicate the orientation (vertical or horizontal) of the line inside the unique shape (the target), a diamond among circles or vice versa. In fixed-feature conditions (top row) the display configuration was fixed within an experimental block, whereas in mixed-feature conditions (bottom row) target and non-target features randomly varied across trials. On a subset of trials, the search display also contained a singleton distractor color. Critically, in no spatial bias blocks (i.e., first half of experiment) this distractor appeared equally often at all search locations, whereas in spatial bias blocks it appeared with a higher probability on one specific location (i.e., high probability distractor location).

To foreshadow our results, while we observed reduced distractor interference at locations with a high distractor probability, counter to many studies, this effect largely reflected intertrial priming. This means that the reported EEG findings from our confirmatory group analyses cannot simply be interpreted in terms of statistical learning. Given the unexpected nature of this finding and given that intertrial priming and statistical learning are naturally conflated (i.e., both contribute to performance in the same direction), we also conducted follow-up analyses at the behavioral level to explore individual differences in sensitivity to location regularities over short versus longer time scales. Based on individual differences we distinguished between observers that did appear to develop spatial expectations and those that did not, to better characterize the observed neural dynamics.

## 1. Methods

Experiment and analyses were based on the OSF preregistration at Open Science Framework (https://osf.io/n35xa/). All data will be made publicly available there upon publication.

A planned number, based on the Wang et al. (2019) study (*N* = 18), of 24 participants (Mean age = 23 years, range 18 – 37; 4 men) participated in the experiment after replacement of 2 participants because EEG preprocessing (for details see below) resulted in exclusion of >30% of trials. Participants were seated in a sound-attenuated and dimly lit laboratory at a viewing distance of ~80 cm.

The paradigm was modeled after the Wang et al. (2019) study. Each trial started with a randomly jittered blank display (0-ms – 350-ms) followed by a 1200-ms fixation display. This display contained a shape that looks like a combination of bulls eye and cross hair, which has been shown to improve stable fixation (Thaler, Schütz, Goodale, & Gegenfurtner, 2013) and that remained visible throughout the trial. Subsequently, a search display appeared with eight shapes in a circular configuration around fixation (radius 4.3°), which remained visible until response. Each display contained one circle (radius 1.05°) and seven diamonds (2.7° x 2.7°), or vice versa, each with a red or green outline on a black background and a horizontal or vertical white line segment (1.6°) in their center. Participants were instructed to keep their eyes at fixation and covertly search for the unique shape in the display and indicate the orientation of the line segment in this target shape via a button press (i.e., ‘z’ or ‘/’ button) as quickly as possible while keeping the number of errors to a minimum. Whereas on 40% of trials all stimuli in the search display had the same color, in the other trials the outline of one of the distractor shapes had a different color than the other stimuli in the display. Participants were instructed to ignore this singleton distractor. Targets and singleton distractors were only presented along the vertical and horizontal axis. Critically, in the first half of the experiment, targets and distractors appeared equally often at these four locations (no spatial bias blocks). In the second half of the experiment^1^, the singleton distractor appeared with 66.7% probability at one of the horizontal locations (counterbalanced across participants), while targets nevertheless continued to appear equally often on all locations (spatial bias blocks) collapsed over distractor-present and -absent trials.

Crucially, in separate experimental sessions (separated by 5-7 days) participants performed the task in the mixed- or the fixed-feature condition (order counterbalanced across participants). The high probability distractor location was kept the same in both sessions. In the mixed-feature condition, targets were equally likely to be a circle among diamonds or a diamond among circles and the singleton distractor colors were also equally likely to be red or green. By contrast, in the fixed-feature condition, a new display configuration was introduced at the start of each block, but target and distractor features (i.e., shape and color) remained fixed throughout a block of trials (features counterbalanced across blocks).

Participants completed 16 blocks of 60 trials per session, each preceded by a sequence of training blocks (60 trials; no spatial bias) until they felt comfortable with the task. At the start of each block participants received written instructions about the dynamics of the upcoming set of trials (e.g., ‘In the upcoming block the target and distractor will have random shapes and colors’). After each block participants received feedback on their performance (i.e., mean reaction time (RT) and accuracy) and were encouraged to take a break. At the end of the second session, a display with only circles, each with a unique number (i.e., 1-8) corresponding to one of the search locations was shown. Participants were asked to indicate (and if necessary, guess) which location they believed had contained the singleton distractor most frequently throughout the experiment.

### Behavioral analysis

All data were preprocessed in a Python environment (Python Software Foundation, https://www.python.org/). As preregistered, behavioral analyses were limited to reaction time (RT) data of correct trials only. RTs were filtered in a two-step trimming procedure: trials with reaction times (RTs) shorter than 200 ms were excluded, after which data were trimmed on the basis of a cutoff value of 2.5 SD from the mean per participant. Remaining RTs were analyzed with repeated measures ANOVAs, where reported p-values are Greenhouse-Geiser corrected in case of sphericity violations, followed by planned comparisons with paired t-tests using JASP software *(JASP-TEAM, 2018)*.

### EEG recording and preprocessing

EEG data were recorded at a sampling rate of 512 Hz using a 64-electrode cap with electrodes placed according to the 10-10 system (Biosemi ActiveTwo system; biosemi.com) and from two earlobes (used as offline reference). As usual for Biosemi, two additional electrodes (common mode sense, CMS, and driven right leg, DRL) were used as reference and ground electrodes during recording. ActiveTwo amplifiers are DC coupled, and signals were stored using ActiView software (Biosemi) with decimation/anti-aliasing filter (5^th^-order Bessel; cutoff frequency, 3-dB attenuation, of 128 Hz) applied to the data streamed to file. Vertical and horizontal EOG (VEOG/HEOG) were recorded via external electrodes placed ~2 cm above and below the eye, and ~1 cm lateral to the external canthi, respectively. EEG from the separate sessions were preprocessed independently (as specified below) after which the two data sets were concatenated.

As preregistered, EEG data were re-referenced off-line to the average of the left and the right earlobe, and subsequently high-pass filtered using a zero-phase ‘firwin’ filter at 0.1 Hz to remove slow drifts. Continuous EEG was then epoched from −1500 – 1100-ms relative to search display onset (to avoid filter artefacts during preprocessing and time-frequency analysis), with trial rejection procedures being limited to smaller time windows (i.e., −1000 – 600-ms). The resulting epochs were baseline normalized during preprocessing using the whole epoch as a baseline to aid detection of noisy electrodes based on visual inspection. Prior to trial rejection, malfunctioning electrodes (*M* = 1, range = 0 - 6) were temporarily removed. EMG contaminated epochs were then identified using an adapted version of an automatic trial-rejection procedure as implemented in the Fieldtrip toolbox (Oostenveld, Fries, Maris, & Schoffelen, 2011). To specifically capture muscle activity we used a 110-140 Hz band-pass filter, and allowed for variable z-score cut-offs per subjects based on within-subject variance of z-scores (de Vries, van Driel, & Olivers, 2017) resulting in an average removal of 9.2% (range 2.3% - 22.4%) of trials thus identified as containing artifacts. Next, ICA as implemented in MNE (method = ‘picard’) was performed on the continuous EEG to identify and remove eye-blink components selectively from the epoched EEG, but not the EOG data. Finally, malfunctioning electrodes were interpolated using spherical splines (Perrin, Pernier, Bertrand, & Echallier, 1989) before the data of the separate sessions was combined.

Throughout the EEG recording, eye movements were monitored using an Eyelink 1000 (SR Research), sampled at 500 Hz, with a remote free-to-move head set-up. Gaze data were analyzed online, so that every time the participant broke fixation, auditory feedback was provided, signaling to the participant to keep their gaze at fixation. In some cases, this auditory feedback was turned off, but only once fixation was relatively stable, as it was considered distracting by a subset of participants when it did occur. The Eyelink data were also analyzed offline. As in our previous work (van Moorselaar & Slagter, 2019), to control for drifts in the eye-tracker, epochs without a saccade (Nyström & Holmqvist, 2010) in the 300 ms pre-display interval were shifted towards fixation. Subsequently, each data sample was converted into a single number specifying the visual deviation from fixation (in visual angle), after which epochs were summarized by a single value indicating the largest deviation in visual degrees from fixation measured in a continuous segment of data of at least 40 ms. Thus, each epoch was summarized by its maximal deviation in visual degrees. The results of the EEG analyses presented next are limited to trials with a fixation deviation <= 1° or in case of missing Eyelink data, trials with sudden sharp jumps in the HEOG as detected via a step algorithm with a window length of 200-ms, a step size of 10 ms, and with a threshold of 20*μV*, which led to the exclusion of 7.7% of the cleaned data (range = 0.7% - 21.8%). As shown in Appendix B this procedure ensured that observed effects could not be explained by systematic eye movements. All EEG preprocessing parameters per subject are summarized at OSF, alongside the raw data files.

### Event-related potential (ERP) analysis

We examined effects of feature- and location-expectations on distractor and target processing using ERPs, which were computed for a prespecified time window from −100 to 600-ms separately per condition of interest. First, epochs were 30 Hz low-pass filtered and baseline corrected using a −100 to 0 ms pre-stimulus baseline period. To enable isolation of lateralized distractor- and target-specific ERP components, ERP analyses focused on trials in which the stimulus of interest (distractor or target) was presented to the left or right of fixation, while the other stimulus was presented on the vertical meridian. Waveforms evoked by the various search displays were collapsed across left and right visual hemifield and left and right electrodes to produce separate waveforms for contralateral and ipsilateral scalp regions. Lateralized difference waveforms were then computed by subtracting the ipsilateral waveform from the corresponding contralateral waveform. In spatial bias blocks, this analysis was limited to trials where the distractor appeared at high probability distractor locations, because there were too few trials with both a distractor at the horizontal low probability location and a target at the vertical midline (*N* = ~30).

Following Wang et al. (2019), we focused the analysis of distractor- and target-elicited Pd and/or N2pc ERP waveforms on electrodes PO7/8. Pd epochs of interest were centered (± 25-ms) around condition-specific positive peaks in distractor and target-tuned waveforms. In many studies, including in Wang et al. (2019), distractor learning is associated with a distractor-evoked early Pd followed by a second positive peak within the N2pc time range (late Pd). As visual inspection of our results suggested that distractor-specific positivities also contained both an early and a late component (Feldmann-Wüstefeld & Vogel, 2019; Feldmann-Wüstefeld, Brandhofer, & Schubö, 2016; Weaver et al., 2017), we searched for positive peaks in a 50 – 150-ms (early Pd) and a 200 – 400-ms (late Pd) in distractor tuned waveforms and only the 50 – 150-ms window in target tuned waveforms. N2pc epochs of interest were centered around condition-specific negative peaks in target-tuned waveforms within a 200 – 400-ms window.

### Time-frequency analysis

We adopted the same analysis approach as reported in Wang et al. (2019). Using Morlet wavelet convolution, EEG time series were decomposed into their time-frequency representation for frequencies ranging from 1 to 40 Hz in 25 logarithmically spaced steps. To create complex Morlet wavelets, for each frequency a sine wave (*e*^*i*2*πft*^, where *i* is the complex operator, *f* is frequency, and *t* is time) was multiplied by a Gaussian (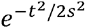, where *s* is the width of the Gaussian). To keep a good trade-off between temporal and frequency precision, the Gaussian width s varied (i.e., *s* = *δ*/(2*πf*), where *δ* represents the number of wavelet cycles, in 25 logarithmically spaced steps between 3 and 12). Frequency-domain convolution was then applied by multiplying the Fast Fourier Transform (FFT) of the EEG data and Morlet wavelets. The resulting signal was converted back to the time domain using the inverse FFT. Time-specific frequency power was defined as the squared magnitude of the complex signal resulting from the convolution.

Condition-averaged lateralized alpha-power (contra- vs. ipsilateral to high probability distractor location) in the pre-stimulus period (i.e., −1000 – 0-ms.) was evaluated via Z-transformed time-frequency power. Z-values were computed by a statistical transformation of real observed lateralized power for each contra- and ipsilateral channel pair with respect to the mean and standard deviation of a “lateralized” power distribution under the null hypothesis of no difference between a contra- an ipsilateral channel pair. This distribution was obtained with a permutation procedure (*N* = 1000), where we swapped electrode pairs for a random subset of trials. This procedure yields a metric that shows interpretable dynamics over time and frequency and has the benefit of not having to select a pre-stimulus interval, the same interval that we are interested in, for baseline normalization.

### Analysis software

Preprocessing and subsequent analyses were performed using custom-written analysis scripts, which were largely based on functionalities implemented within MNE (Gramfort et al., 2014). These scripts can be downloaded at https://github.com/dvanmoorselaar/DvM.

## 2. Results

### Search times

Exclusion of incorrect responses (6.5%) and data trimming (3.1%) resulted in an overall loss of 9.6% of the behavioral data. We first analyzed distractor costs as a function of search condition (Figure 2A/B). A repeated measures ANOVA with the within subject factors Condition (mixed-features, fixed features) and Distractor (present, absent) using only data from no spatial bias blocks showed faster RTs in fixed- than in mixed-feature conditions (main effect Condition: *F* (1, 23) = 31.5, *p* < 0.001, 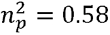) and reliable distractor interference (main effect Distractor: *F* (1, 23) = 119.6, *p* < 0.001, 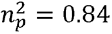). Importantly, a Condition by Distractor interaction (*F* (1, 23) = 20.6, *p* < 0.001, 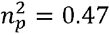) showed that distractor interference was larger in the absence of feature foreknowledge, in the mixed-feature condition (862 vs. 937 ms; *t* (23) = 7.4, *p* < 0.001, *d* = 1.52), compared to when features were entirely predictable, in the fixed-feature condition (675 vs. 694 ms; *t* (23) = 5.6, *p* < 0.001, *d* = 1.14). Notably, while distractor interference was reduced, it was still far from being completely absent in the fixed-feature condition. Conceivably, this may have been the case because in each block observers were introduced to a new display configuration so that target and distractor features, while fixed within a given block, hence varied from block-to-block. Indeed, an exploratory running regression analysis (Figure 2B) examining distractor costs, in which trials per experimental block were grouped in bins of 10 trials, yielded an averaged negative slope of −4.8 ms over bins in the fixed-feature condition that significantly deviated from zero (*t* (23) = 2.2, *p* = 0.039), whereas that slope (*M* = 1.0) was non-significant in the mixed-feature condition (*t* (23) = 0.5, *p* = 0.63). Pairwise comparisons confirmed that distractor costs were reliable across all bins in mixed-feature condition (all *t*’s > 4.0, all *p*’s < 0.001, all *d*’s > 0.082), but only in the first two bins in fixed-feature condition (all *t*’s > 2.7, all *p*’s < 0.0013, all *d*’s > 0.55) and not in bins 3 – 5 (all t’s < 2.0, all p’s > 0.057, all *d*’s < 0.41). This finding is in line with the notion that distractor inhibition critically relies on previous experiences and that to resist distractor interference in fixed-feature conditions, a specific target template needs to be accompanied by expectation-dependent tuning to distractor features (Vatterott & Vecera, 2012).

**Figure 2.**
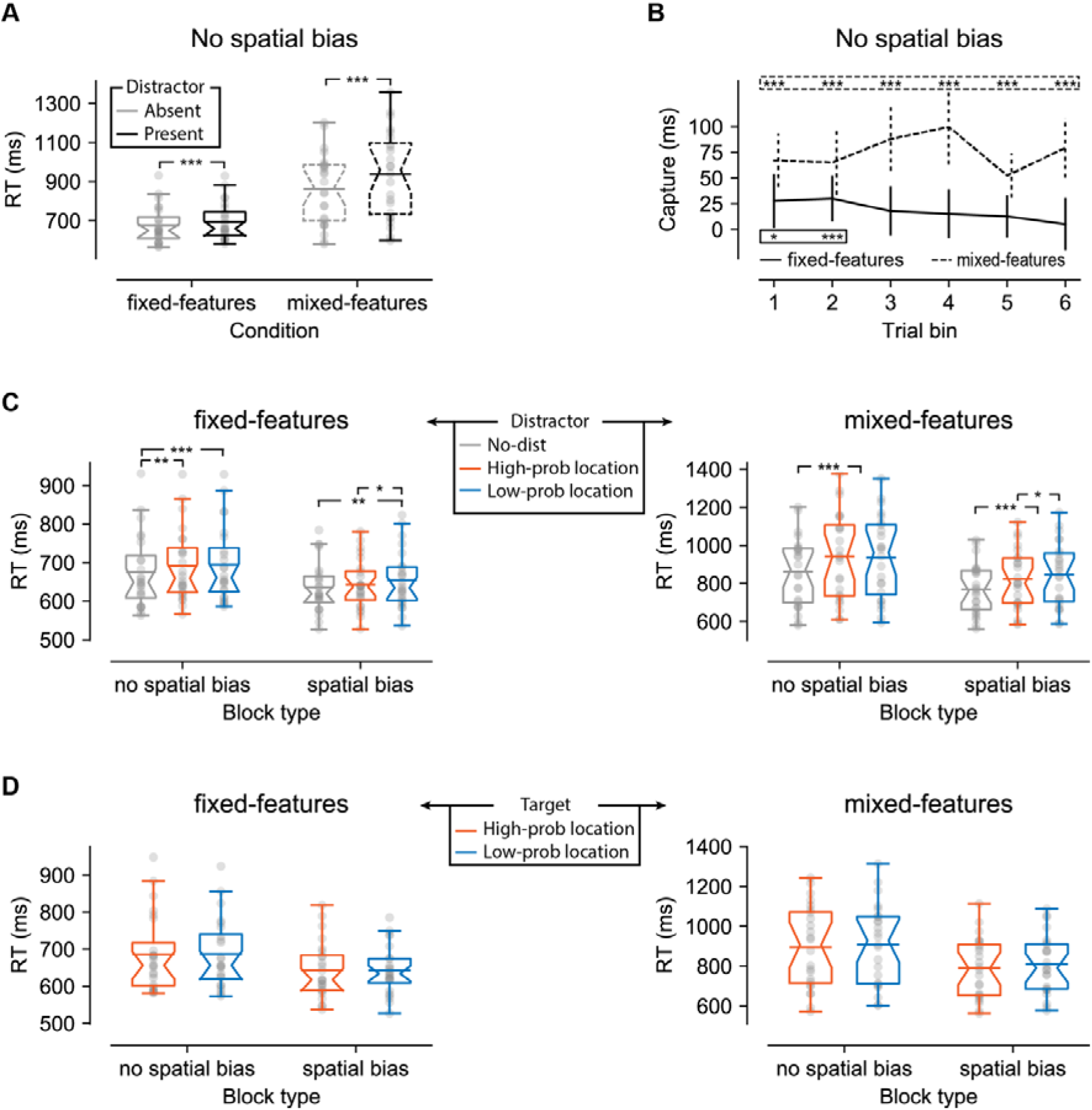
Behavioral findings visualized by notched boxplots with solid horizontal lines corresponding to the mean. (**A**) Reaction times in no spatial bias blocks shown separately for distractor absent and distractor present trials in the fixed-features and mixed-features conditions. (**B**) Binned distractor interference (distractor present – distractor absent RT) per conditions. Each trial bin corresponds to 10 trials (bin 1: trial 1 – trial 10, bin 2: trial 11 – trial 20, etc). Error bars represent condition-specific, within-subject 95% confidence intervals (Morey, 2008) (**C**) Reaction times as a function of distractor location in no spatial bias and spatial bias blocks for fixed-features (left) and mixed feature (right) conditions. Note that in no spatial bias blocks, distractor locations were artificially coded as high and low probable distractor locations. (**D**) Reaction times as a function of target location in no spatial bias and spatial bias blocks for fixed-features (left) and mixed feature (right) conditions. Data is based on distractor absent trials only. The bars in the boxplots and all subsequent boxplots are whiskers, which extend to the most extreme, non-outlier points.

We next determined how distractor interference was modulated when in addition distractor location learning was possible. To this end, we entered RTs into a repeated measures ANOVA with the within subjects’ factors Block Type (no spatial bias, spatial bias), Condition (mixed-features, fixed-features) and Distractor Location (high probability distractor location, low probability distractor location). For this purpose, within no spatial bias blocks, distractor locations were artificially coded as ‘high’ and ‘low’ probable locations as the spatial bias was only introduced in the second half of the experiment. Overall RTs were faster in spatial bias blocks (main effect of Block Type: *F* (1, 23) = 62.7, *p* < 0.001, 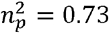) and in the fixed-feature condition (main effect of Condition: *F* (1, 23) = 38.6, *p* < 0.001, 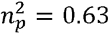). Crucially, as visualized in Figure 2C, and reflected in a Block Type by Distractor Location interaction (*F* (1, 23) = 14.2, *p* < 0.001, 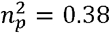), distractors were more efficiently ignored at high probability distractor locations in spatial bias blocks, both in mixed- (*t* (23) = 2.1, *p* = 0.047, *d* = 0.43) and fixed-feature conditions (*t* (23) = 2.3, *p* = 0.028, *d* = 0.48), yet no such benefit was observed in no spatial bias blocks (all *t*’s < 1.0, all *p*’s > 0.33, all *d*’s < 0.20). Importantly, despite the spatial bias distractors continued to produce reliable interference (i.e., relative to no distractor trials) at both the low and high probability locations in mixed-feature blocks (all *t*’s > 6.5, all *p*’s < 0.001, all *d*’s > 1.33) and in fixed-feature blocks, although while this effect was robust at low probability locations (t (1,23) = 3.3, *p* = 0.003, *d* = 0.68), it was only marginally significant at high probability distractor locations (t (1,23) = 2, *p* = 0.055, *d* = 0.41). Together these findings may suggest that both feature- and location-based distractor expectations reduced distractor interference, and may interact in distractor inhibition.

Arguably, a significant proportion of the observed RT benefits in the preceding analysis are due to short-lasting, intertrial location priming (Maljkovic & Nakayama, 1994). A standard control in the literature to establish whether the suppression attributed to expectations derived from past experience or statistical learning is independent from inter-trial (priming) effects, is to repeat the same analysis after eliminating all trials where the distractor location repeated, although it should be noted that elimination of such immediate repetitions still allows for priming effects resulting from more distant trials in the past (Maljkovic & Nakayama, 1994). Using this control, previous studies have shown that the effect of distractor location likelihood remains significant, which has been taken as evidence that distractor probability effects cannot solely be explained in terms of intertrial location priming (e.g., (Failing, Feldmann-Wustefeld, et al., 2019; Ferrante et al., 2018; Sauter et al., 2018). Here, however, while numerically distractors were still ignored more efficiently in spatial bias blocks at high probability locations, the Block Type by Distractor Location interaction was no longer significant (*F* (1, 23) = 1.8, *p* = 0.19, 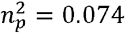). This unexpected finding may indicate a large contribution of intertrial priming and suggests that statistical distractor location learning, if at all present, was less robust than in other studies using similar paradigms.

However, intertrial repetition effects may not only reflect intertrial priming (Feldmann-Wüstefeld & Schubö, 2016; van Moorselaar & Slagter, 2019; Won, Kosoyan, & Geng, 2019). Specifically, statistical learning can further increase the effect of intertrial priming and by subtracting out the intertrial priming effect, statistical learning effects are also eliminated. Indeed, an explorative analysis limited to intertrial priming trials only showed that distractor costs were significantly reduced at high probability distractor locations in spatial bias blocks, but not at low probability distractor locations, although these results should be interpreted with much caution given the low number of observations at low probability distractor locations in spatial bias blocks (*N* = ~4.5). Therefore, in an exploratory analysis, limited to distractor present trials in spatial bias blocks, we adopted a linear mixed models’ approach (for details see caption Table 1)(Bates, Mächler, Bolker, & Walker, 2014). In contrast to conventional ANOVA approaches, in linear mixed models, data is not averaged but grouped per participant. Critically, a range of continuous and categorical variables can be added to a single model such that rather than excluding a large subset, which inevitably reduces power (Brysbaert & Stevens, 2018), intertrial transitions can be included as a control factor allowing for a more refined control of intertrial priming (i.e., all intertrial transitions, D_n-1_ – D_n_, D_n-1_ – T_n_, T_n-1_ – D_n_, T_n-1_ – T_n_, that potentially modulate the statistical learning effect; see supplementary material of Sauter et al., 2018 for an extensive discussion). As shown in Table 1, this analysis showed that while both distractor (D-D) and target (T-T) repetitions contributed significantly, overall RTs were reliably faster at high probability distractor locations after controlling for such intertrial repetitions. Together these findings demonstrate that while the observed spatial probability effects to a large extent reflected intertrial repetition, in contexts where the distractor location was more frequently repeated, regularities across multiple trials also contributed to performance. Confirmatory studies are necessary to replicate these exploratory findings.

**Table 1.** Estimates for mixed effects model, using Satterthwaite’s method for approximating degrees of freedom (Luke, 2017). Following guidelines by (Barr, Levy, Scheepers, & Tily, 2013), the model had a participant grouping variable, with a random effect structure including an intercept, Distractor Location, and Target Location and as fixed variables Distractor location, Target location, Condition, and the control variables indexing four different forms of intertrial priming (D_n-1_ – D_n_, D_n-1_ – T_n_, T_n-1_ – D_n_, T_n-1_ – T_n_). The table shows the unstandardized estimates (*β*), the standard error (SE), estimated degrees of freedom (df) and the corresponding t and p values. We used sum coding (−1 vs 1) for all control factors in the model and dummy coding (0 vs 1) for all other factors in the model, to facilitate interpretation of the statistics (see baselines within the table).

| | $\beta$ | SE | df | t | p |
| --- | --- | --- | --- | --- | --- |
| Condition (baseline = mixed) | -183.0 | 26.6 | 24.0 | -6.8 | <0.001 |
| Distractor location (baseline = low probability)<sup>2</sup> | -17.0 | 5.8 | 27.8 | -2.9 | 0.007 |
| Target location (baseline = low probability) | -11.8 | 10.4 | 25.4 | -1.1 | 0.27 |
| $D_{n-1} - D_n$ | -21.2 | 4.4 | 12478.7 | -4.8 | <0.001 |
| $D_{n-1} - T_n$ | 6.2 | 4.4 | 12495.1 | 1.4 | 0.16 |
| $T_{n-1} - D_n$ | 8.2 | 5.9 | 12500.0 | 1.4 | 0.16 |
| $T_{n-1} - T_n$ | 46.4 | 4.3 | 12500.8 | 10.8 | <0.001 |

To further examine the influence of distractor location regularities on spatial suppression, we repeated the preceding analyses, but now tuned to target locations and only including distractor absent trials (Figure 2D). Counter to previous findings (e.g., (Failing, Feldmann-Wustefeld, et al., 2019; Wang et al., 2019), but consistent with the exploratory linear mixed models analysis (Table 1), there was no evidence that target processing was impaired at high probability distractor locations across conditions (Block Type by Distractor Location interaction: *F* (1, 23) = 0.048, *p* = 0.83, 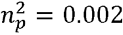), not even when the analysis was limited to the mixed-feature condition (t = 0.8, *p* = 0.44, *d* = 0.16), where we a priori did expect impaired target processing.

Finally, we examined if participants could correctly identify the high probability distractor location. Eight out of 24 participants correctly identified the high probability distractor location (chance level = 25% or 6 participants; assuming that participants noticed that stimuli only appeared on cardinal axes) suggesting that the spatial bias may have been noted by at least a subset of participants. However, the average deviation from the high probability location was 2.1 and thus not above chance (chance level = 2). After excluding these eight participants, the results from the distractor and target location tuned ANOVA’s did not change (i.e., same effects were reliable), but the effect of Distractor location (i.e., the statistical learning effect) in the exploratory linear mixed models was no longer significant (*p* = 0.14). This may suggest that explicit recognition of the high probability distractor location strategically enhanced the observed statistical learning effect. Note however, that this analysis inevitably had less observations and results should thus be interpreted cautiously, also since simply on the basis of chance, we can expect 6 subjects to pick the high probability distractor location even when unaware.

### 3.1 Interim discussion and exploratory analyses

Consistent with previous studies (e.g., (Bacon & Egeth, 1994; Theeuwes, 1992), in mixed-feature blocks the colored distractor singleton reliably interfered with attentional selection, while in fixed-feature blocks this interference was reduced. That is, while interference was prevalent early on in a fixed-feature block when observers were introduced to a new stimulus configuration, it disappeared once they gained more experience with the distracting information. These findings are in line with the idea that learned expectations play a critical role in distractor inhibition (van Moorselaar & Slagter, 2020). Also replicating previous findings, introducing a spatial bias greatly reduced distractor interference at high probability distractor locations. Yet, our findings unexpectedly differ from previous findings in two important aspects. First, distractor suppression was not accompanied by impaired target processing at high probability locations (Wang & Theeuwes, 2018b). Second, whereas previous studies have shown that spatial distractor suppression cannot solely be explained in terms of intertrial location priming (Failing, Feldmann-Wustefeld, et al., 2019; Ferrante et al., 2018), here distractor suppression at high probability locations appeared to be largely driven by lingering biases from the preceding trial. These two discrepancies are especially relevant as we aimed to investigate whether neural mechanisms underlying learning from spatial regularities differ under conditions that either allow for the development of feature expectations or not.

One notable difference with previous studies (Wang & Theeuwes, 2018b; Wang et al., 2019) is that in the currently study, the high probability location was not introduced immediately, but only halfway through the experiment. It has been shown that different strategies may be applied to the search task depending on the specifics of the individual paradigms (Zhang et al., 2019). Although speculative, in the current study, observers may generally have been less prone to learn more subtle probabilities once they already had developed a strategy specifically tuned to conditions without such probabilities. If distractor rejection is efficient in the absence of spatial regularities, there is also less need for spatial distractor learning, simply because distractors are already efficiently ignored. Indeed, an explorative correlational analysis between distractor interference in the first half of the experiment (i.e., distractor present vs. absent) and selection history effects (distractor at high vs. low probability location) yielded a robust positive correlation both in mixed- (*r_s_* = 0.43, *p* = 0.038) and fixed-feature blocks (*r_s_* = 0.45, *p* = 0.026). That is, participants who exhibited high distractor interference when its location could not be predicted in advance in the first half of the experiment, generally were more sensitive to the high probability distractor manipulation in the second half of the experiment, consistent with the idea that the magnitude of selection history biases depends on the amount of distractor interference (Failing & Theeuwes, 2020; Sauter et al., 2018). While this correlation appears to be in line with the idea that statistical learning becomes increasingly sensitive to individual differences in initial task performance when the spatial regularity is introduced at a later stage, it should be noted that same correlation was no longer significant after controlling for intertrial distractor priming (all *r_s_*‘s > 0.18, all *p*‘s < 0.38).

To nevertheless explore these potential individual differences further we post-hoc divided participants in two groups (‘learners’ and ‘non-learners’) based on whether or not target processing was impaired at high probability distractor locations. As a priori, we expected this impaired target processing to be specific to mixed-feature conditions, we only used this condition to label individual participants as ‘learners’ and ‘non-learners’ on the basis of whether the target impairment score (i.e., target at high probability distractor location vs. target at low probability distractor location) was larger or equal to/smaller than zero, respectively^3^. The logic here is that if this division is not simply taking two halves of a normal distribution around zero, but is truly diagnostic of individual differences, then spatial suppression should not only also go above and beyond intertrial repetition in ‘learners’, but one would also expect neural markers of suppression to differ between groups (e.g., a Pd to targets at high probability distractor location in spatial bias blocks in the mixed-features condition in the learners, but not in the non-learners). Note that importantly, distinctions between ‘learners’ and ‘non-learners’ could not be explained by the order in which they completed both EEG sessions (i.e., 5 and 7 learners identified in session 1 and 2, respectively).

Intriguingly, the two groups, which were thus selected on the basis of a target impairment score in spatial bias blocks (i.e., second half of experiment), already differed in no spatial bias blocks (i.e., first half of experiment), with ‘learners’ exhibiting larger distractor costs than ‘non-learners’ (Distractor by group interaction: *F* (1, 22) = 5.1, *p* = 0.034, 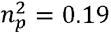), while being, although only marginally significant, slower overall (main effect Group: *F* (1, 22) = 4.0, *p* = 0.059, 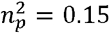). More importantly, in spatial bias blocks, a Distractor Location by Group interaction (*F* (1, 22) = 5.7, *p* = 0.026, 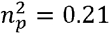) showed that ‘learners’ (high vs low probability distractor location fixed features: *M* RT= 654.5 ± *SD* = 69.6 vs 672.0 ± *SD* = 81.0; high vs low probability distractor location mixed features: *M* RT = 872.8 ± *SD* = 147.1 vs 917.5 ± *SD* = 146.2) were less distracted by distractors at high vs. low probability distractor locations than ‘non-learners’ (high vs low probability distractor location fixed features: *M* RT = 631.8 ± *SD* = 60.0 vs. 637.2 ± *SD* = 67.4; high vs low probability distractor location mixed features: *M* RT = 772.0 ± *SD* = 152.1 vs. 773.0 ± *SD* = 160.6). Moreover, the same interaction remained reliable after exclusion of trials in which the distractor location repeated (*F* (1, 22) = 5.6, *p* = 0.027, 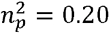), demonstrating that in ‘learners’ distractor suppression could not be explained by intertrial priming alone (*F* (1, 11) = 9.0, *p* = 0.012, 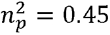). Finally, although ‘learners’ were selected on the basis of the target impairment score in the mixed-feature condition (‘learners’ vs. ‘non-learners: *M* = 33.6 ± *SD* = 11.7 vs. −50.7 ± *SD* = 44.8), they also exhibited, although only marginally significant (*t* (11) = 2.0, *p* = 0.07, *d* = 0.57), impaired target processing at high probability distractor location in the fixed-feature condition (‘learners’ vs. ‘non-learners: *M* = 23.0 ± *SD* = 39.9 vs. −18.5 ± *SD* = 39.7). Together, these findings suggest that our group division was not an arbitrary split, but rather separated participants that were sensitive to the introduced statistical regularity over longer time scales from those where suppression to a larger extent was driven by intertrial repetition. Therefore, in addition to the preregistered whole group ERP and time-frequency analyses, we also separate between ‘learners’ and ‘non-learners’ in a set of exploratory analyses.

### 2.2 ERPs

We next examined effects of distractor feature and location regularities on the lateralized Pd and N2pc ERP components. Visual inspection of Figure 3 shows that in the no spatial bias blocks, a positive difference, corresponding to the Pd, emerged to lateral distractors with midline targets. Notably, while this positivity consisted of an early and a late component in the fixed-feature condition, in the mixed-feature condition, in which participants could not predict what the distractor would look like in advance, only a late Pd can be observed. By contrast, in the spatial bias blocks, at the high probability distractor location, the distractor Pd was virtually identical in mixed- and fixed-feature conditions.

**Figure 3.**
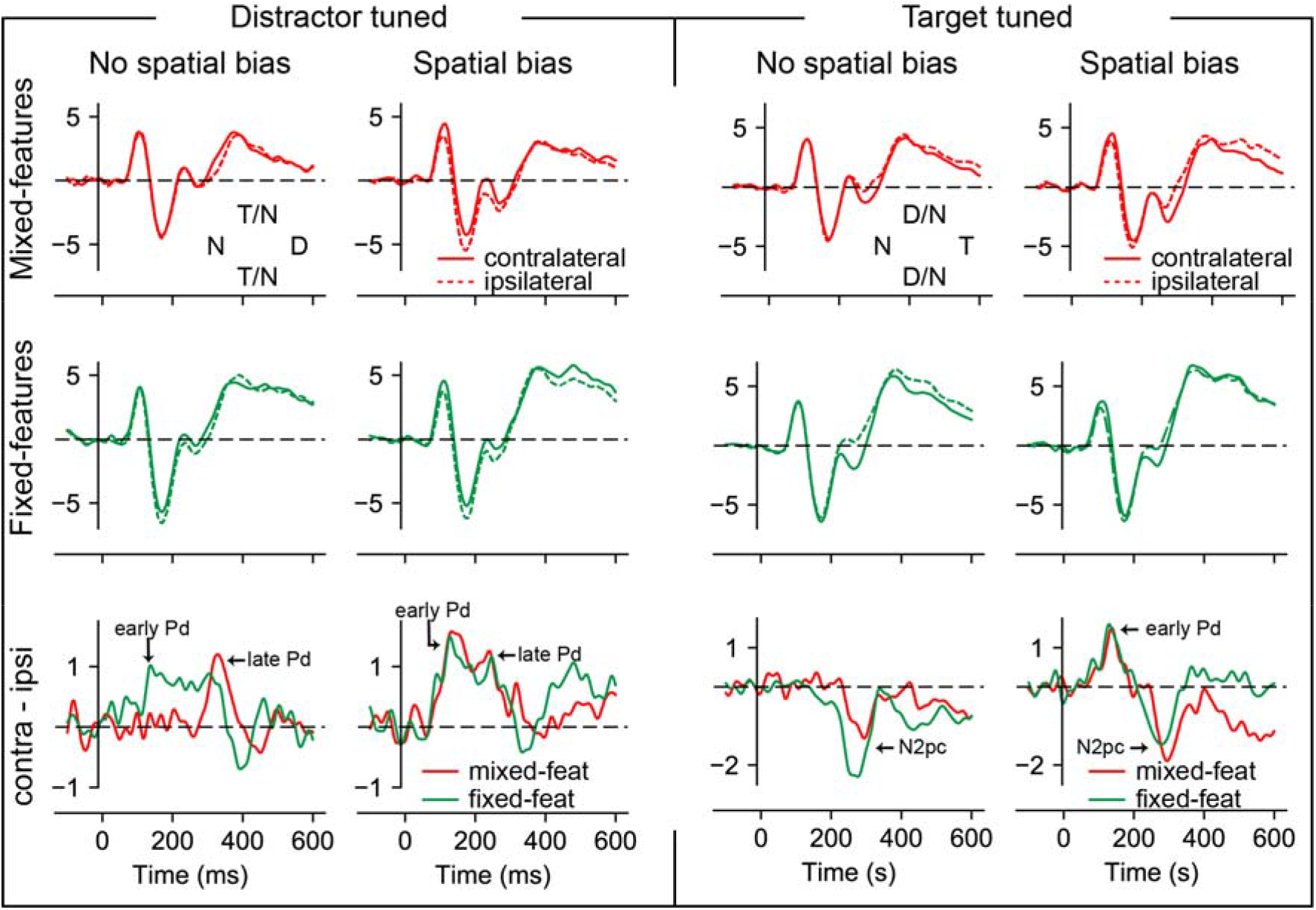
Electrophysiological results from search trials with lateral distractors (distractor tuned ERPs; columns 1 and 2) and with lateral targets (target tuned ERPS; columns 3 and 4), separately for no spatial bias (columns 1 and 3) and spatial bias blocks (columns 2 and 4). Rows 1 and 2 show the contra- and ipsilateral waveforms for the mixed- and fixed-features conditions, respectively. Row 3 shows the difference between contra- and ipsilateral waveforms for the mixed- and fixed-features conditions. Microvolts are plotted on the y-axes. Inserts shows the schematic target (T), distractor (D) and non-target N) positions.

Differences in contra- vs. ipsilateral voltage elicited by lateralized distractors at “high” probability distractor locations at electrodes PO7/8 were therefore analyzed separately in the early and late Pd windows. The early Pd peaked on average in no spatial bias blocks at 100 ms (mixed-feature) and at 137 ms (fixed-feature), and in spatial bias at 133 ms (mixed-feature) and at 129 ms (fixed-feature). A repeated measures ANOVA with peak amplitude in a 50 ms window surrounding the peak as the dependent variable and Condition (mixed features, fixed features), Block Type (no spatial bias, spatial bias) and Hemifield (contralateral, ipsilateral) revealed that the main effect of Hemifield (*F* (1, 23) = 4.5, *p* = 0.045, 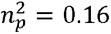), was accompanied by a significant three-way interaction (*F* (1, 23) = 8.1, *p* = 0.009, 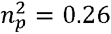). This confirmed that the early Pd was reliably larger in fixed-than mixed-feature conditions, but only in no spatial bias blocks (*t* (23) = 3.7, *p* = 0.001, *d* = 0.75), and not at high probability distractor locations in spatial bias blocks (*t* (23) = 0.1, *p* = 0.93, *d* = 0.02). The late Pd peaked on average slightly later in no spatial bias blocks at 328 ms (mixed-feature) and at 305 ms (fixed-feature), than in spatial bias at 240 ms (mixed-feature) and at 246 ms (fixed-feature), but did not differ in amplitude across conditions. This was statistically reflected in the fact that in the late Pd analysis, the main effect of Hemifield (*F* (1, 23) = 4.4, *p* = 0.047, 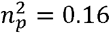) was no longer accompanied by any interactions (all *F*’s < 0.7, all *p*’s > 0.40). Moreover, an exploratory peak latency analysis^4^ within the late Pd window showed significant main effects of Condition and Block Type (all *F*’s > 4.7, all *p*’s < 0.041 all 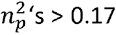), reflecting an earlier late Pd as a function of feature and spatial regularities, respectively. In addition, this analysis revealed a Condition by Block Type interaction (*F* (1, 23) = 5.6, *p* = 0.027, 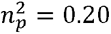), which reflected that fact that whereas the late Pd peaked later in mixed-features (*M* = 314-ms) than in fixed-features (*M* = 277-ms) conditions (*t* (23) = 2.7, *p* = 0.012, *d* = 0.56) in no spatial bias blocks, this difference between mixed-features (*M* = 269-ms) and fixed-features (*M* = 265-ms) conditions disappeared in spatial bias blocks (*t* (23) = 0.4, *p* = 0.67, *d* = 0.088). Together, these results suggest that an early suppressive mechanism, as reflected in the early Pd, can only develop in mixed-feature conditions as a function of spatial regularity, and that a later stage suppressive mechanism, as captured by the late Pd, onsets earlier in time when regularities occur either at the spatial or the feature level.

The presence of a Pd in fixed-feature conditions irrespective of the spatial bias suggests that the observed reduction in distractor interference in the fixed-feature condition cannot solely be explained by a more precise target template, but may also reflect feature-based experience-dependent distractor inhibition. In line with this, behaviorally, in fixed feature conditions, distractor interference gradually decreased over the course of a block as observers increasingly gained experience with the distracting information (Figure 2B). One would hence expect that in no spatial bias blocks, the Pd also emerged gradually. To test this prediction, following the behavioral analysis where distractor interference gradually decreased within each block, in an explorative analysis, we analyzed the signed area under the curve (i.e., the positive area; baseline is zero) in a 100 – 400-ms window using only non-spatial bias blocks with a repeated measures ANOVA with the within subjects’ factors Condition (mixed-features, fixed-features) and Trial Bin (1 – 20, 21 – 60). Here, we focused on area under the curve rather than mean amplitude as this method is less sensitive to potential latency differences between individuals. A reliable interaction (*F* (1, 23) = 5.5, *p* = 0.028, 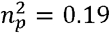) demonstrated that in fixed-feature blocks the Pd area was indeed larger in the final 40 trials of the block then in the first 20 trials (*t* (23) = 2.4, *p* = 0.027, *d* = 0.48), whereas no such difference was observed in mixed-feature conditions (*t* (23)= 1.3, *p* = 0.21, *d* = 0.27) (Figure 4). Intriguingly, while the same feature-based learning effect was absent in spatial bias blocks (*F* (1,23) = 1.2, *p* = 0.29, 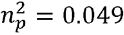), in fixed-feature conditions the Pd area was larger in spatial bias blocks than in non-spatial blocks, not only for the final 40 (*t* (23) = 2.5, *p* = 0.019, *d* = 0.51), but already for the first 20 trials in a block (*t* (23) = 3.4, *p* = 0.003, *d* = 0.69). Together these findings suggest that the inhibitory mechanism reflected by the Pd was dependent on both feature-based and location-based information in the fixed feature conditions, whereas in mixed-feature conditions, distractor suppression was purely spatially informed.

**Figure 4.**
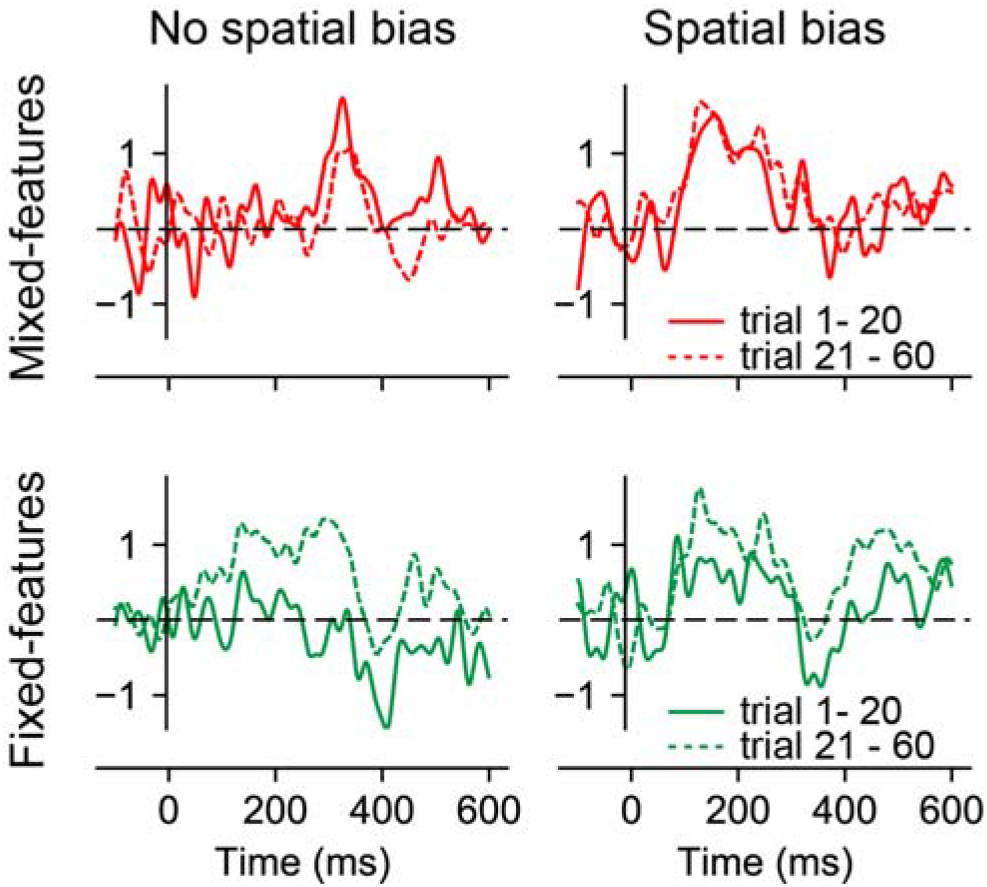
In fixed features conditions in no spatial bias blocks the Pd only emerged after some experience was gained with the distracting information, suggestive of feature-based learning. Results from exploratory analysis: difference between contra- and ipsilateral waveforms across conditions separately for first 20 (solid lines) and last 40 trials (dashed lines) within a block. Microvolts are plotted on the y-axes.

We next examined effects of feature and spatial regularities on target processing. Lateralized ERPs evoked by lateral targets with midline (or absent) distractors are shown in Figure 3. As can be seen, targets evoked a clear N2pc in all conditions. In no spatial bias blocks, this N2pc was larger in amplitude in fixed-feature than in mixed-feature conditions. As expected based on Wang et al. (2019), after introduction of the spatial bias, targets at high probability distractor locations first appeared to elicit an early Pd, both in the fixed-features and mixed-features condition. Yet, counter to Wang et al. (2019), this early Pd was immediately followed by an N2pc (rather than a late Pd in their study).

Voltages elicited by lateralized targets within the N2pc interval (295 ± 25-ms for no spatial bias/mixed features; 275 ± 25-ms for no spatial bias/fixed features; 295 ± 25-ms for spatial bias/ mixed features; 279 ± 25-ms for spatial bias/fixed features) were analyzed as above. While the N2pc was reliable across all conditions (main effect Hemifield: *F* (1, 23) = 9.5, *p* = 0.005, 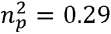), there was no evidence that it differed between conditions or was reduced in amplitude at high probability distractor locations, as neither Condition nor Block Type interacted with Hemifield (all *F*’s < 1.9, all *p*’s > 0.18). At the same time, despite being clearly visible in the group-average lateralized waveforms (Figure 3), an analysis of the early Pd interval did not yield a Block Type by Hemifield interaction (*F* (1,23) = 1.5, *p* = 0.24, 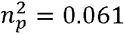), nor did planned-pairwise comparisons identify hemispheric differences in any of the conditions (all *t*’s < 1.5, all *p*’s > 0.15). This suggests that at a group level the early Pd at high probability distractor locations was not reliably evoked by targets across subjects (but see below).

One potential explanation for the finding that the target-evoked Pd at high probability distractor locations was not reliable, is that it is selectively elicited by a subset of participants. Note that our exploratory behavioral analysis suggested that some participants demonstrated typical markers of spatial statistical learning (including impaired target processing), while in other participants effects largely reflected intertrial repetition. Therefore, in an exploratory analysis using the condition specific time windows as in the preceding analyses, we included Group (‘learners’, ‘non-learners’) as a between subjects’ factor to explore whether the early target-evoked Pd was selectively elicited by the learners. Surprisingly, however, a Group by Block Type interaction (*F* (1, 22) = 5.2, *p* = 0.032, 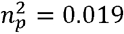) and a main effect of Group (*F* (1, 22) = 4.8, *p* = 0.039, 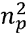 = 0.018) showed that ‘non-learners’ rather than ‘learners’ were driving the target Pd effect (see Figure 5), strongly suggesting that the early Pd at the high probability distractor location discussed above thus largely reflects intertrial priming, as some studies have also suggested (Gokce, Geyer, Finke, Müller, & Töllner, 2014), instead of spatial statistical learning. Consistent with this, in spatial bias blocks the early Pd in response to distractors was virtually absent in ‘learners’, but especially pronounced in ‘non-learners’ (main effect of Group: *F* (1, 22) = 4.2, *p* = 0.053, 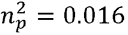). Intriguingly, the opposite pattern was observed for the late Pd, which appeared largely driven by learners, although this effect was not statistically significant (*F* (1,22) = 0.18, *p* = 0.67, 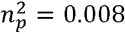). This apparent dissociation may suggest that early and late distractor Pd’s observed at the group level in spatial bias blocks did not reflect two sequential suppressive processes, but instead individual differences in early and late suppression mechanisms. Moreover, the current results may suggest that the early Pd is especially sensitive to intertrial priming.

**Figure 5.**
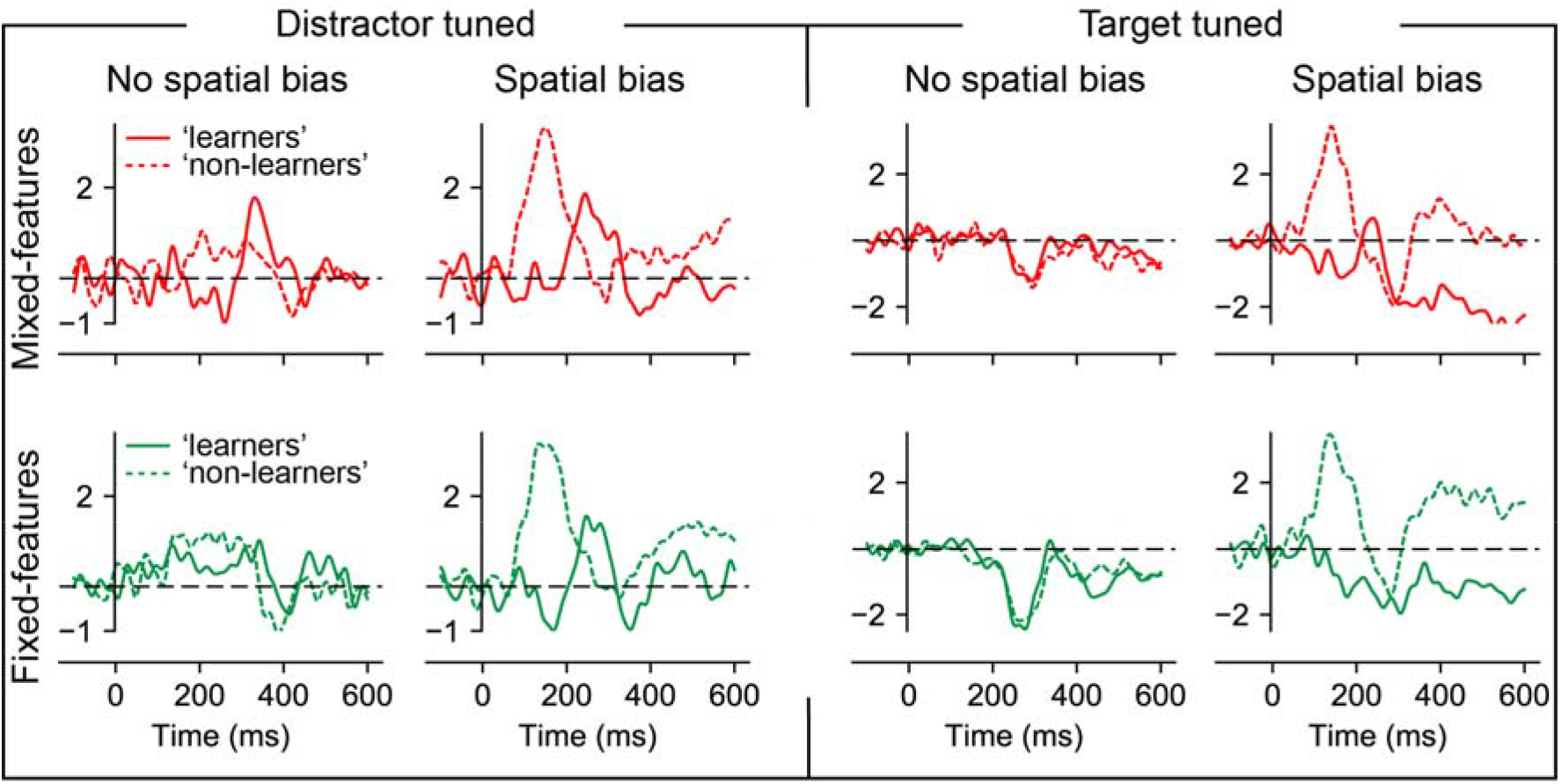
Individual differences in spatial statistical learning were associated with individual differences in the early and late Pd. Results from exploratory analysis: difference between contra- and ipsilateral waveforms across conditions separately for ‘learners’ and ‘non-learners’. Microvolts are plotted on the y-axes.

### 2.3 Time-frequency analysis

To investigate whether the introduction of the high probability distractor location lead to anticipatory spatial inhibition, we focused our analysis on pre-stimulus alpha-band oscillatory activity (7.5 – 12 Hz; Figure 6A/B). Counter to Wang et al. (2019), after introduction of the spatial bias, no reliable increase in pre-stimulus alpha-band power was observed at contralateral compared to ipsilateral electrodes relative to the high probability distractor location in the mixed-feature condition, nor in the fixed-feature condition. That is, cluster-based permutation tests across time and alpha frequencies between spatial and non-spatial blocks separately for the fixed-features and mixed-features conditions did not yield any reliable differences, neither did the same tests across time on averaged alpha power. Also, a Condition (mixed-features, fixed-features) by Block Type (no spatial bias, spatial bias) repeated measures ANOVA on averaged alpha power within the anticipatory window (−1000 – 0ms) did not show reliable effects (all *F*’s < 2.0, all *p*’s > 0.17).

**Figure 6.**
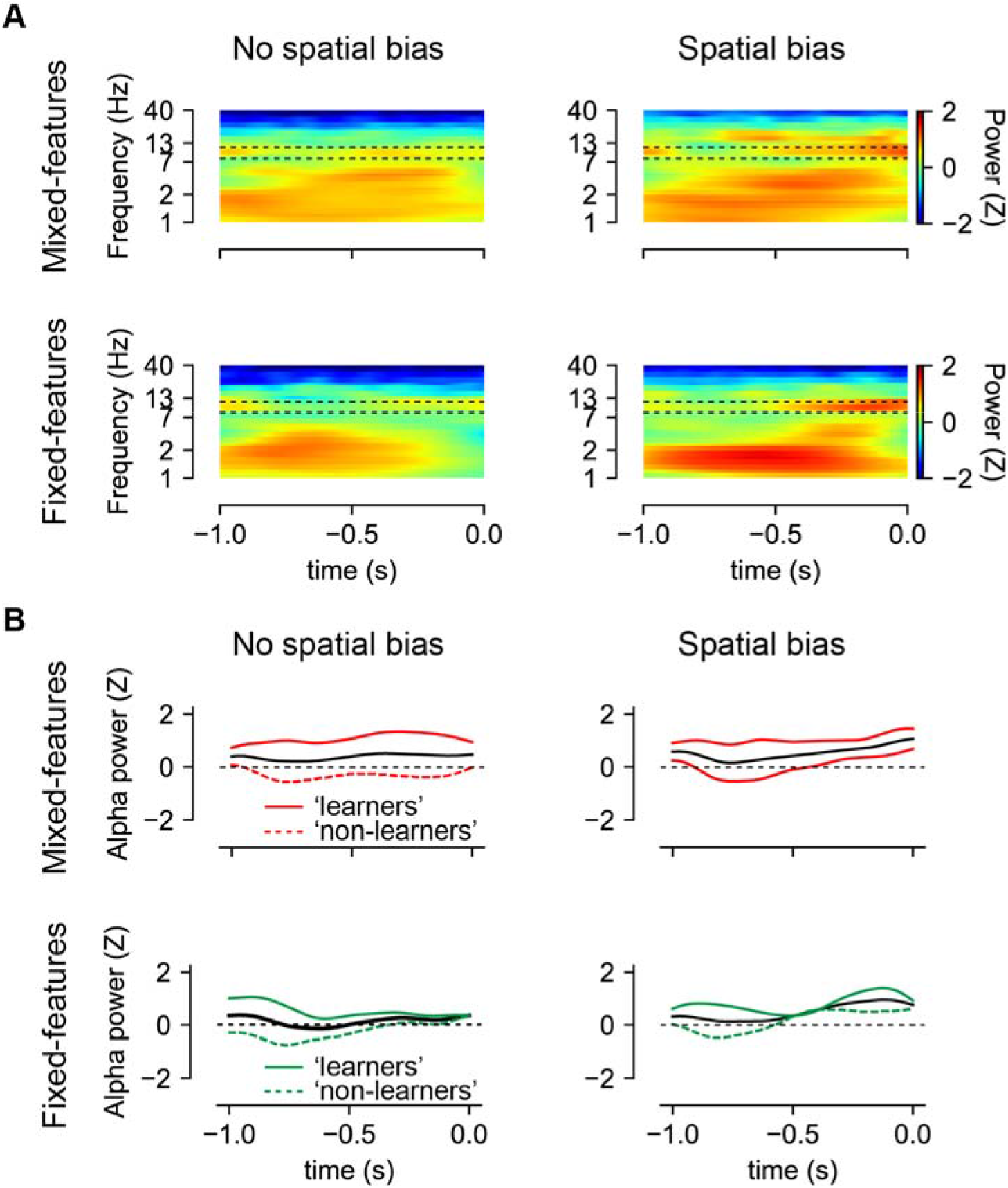
Distractor location learning was not associated with any changes in pre-stimulus alpha-band activity contralateral to the likely distractor location. Shown is posterior lateralized Z scored power. (**A**) Time-frequency representation of contralateral minus ipsilateral (relative to the high probability distractor location) power averaged over electrodes O1/2, PO3/4 and PO7/8. Dotted black outlines show the frequency bands of interest (7.5 – 12 Hz) (**B**) Time series of contralateral minus ipsilateral averaged alpha power (7.5 – 12 Hz) averaged over electrodes O1/2, PO3/4 and PO7/8. Solid black lines show the group mean, whereas colored lines show the effect separately for participants labeled as ‘learners’ and ‘non-learners’.

It is possible that the development of generic suppression by alpha-band activity was not observed because the spatial bias effect largely reflected intertrial repetition rather than learning across longer time scales. In an exploratory analysis we therefore next investigated whether lateralized alpha power modulations were specific to participants who exhibited slower search times when the target was presented at the high probability distractor location (i.e., the learners). As visualized in Figure 6B, however, here again there was no evidence that lateralized alpha power was increased in spatial bias blocks relative to no spatial bias blocks, neither in ‘learners’ nor in ‘non-learners’ (all Group interactions: all *F*’s < 1.0, all *p*’s > 0.33). While these effects should be interpreted cautiously, the absence of anticipatory alpha modulations is in line with findings from two other EEG studies (Noonan et al., 2016; van Moorselaar & Slagter, 2019) that showed clear expectation-based distractor suppression and thus argue against the idea that activity of brain regions representing the high probability distractor location are suppressed in advance, at least as indexed by modulations of alpha-band activity.

## 3. General discussion

The aim of the present study was to gain more insight into the neural mechanisms underlying distractor inhibition as a function of whether it’s specific features and/or likely location could be predicted in advance or not. To this end, observers searched for a unique shape among homogeneous non-target shapes, either in conditions where target and non-target shapes randomly swapped or were fixed (and hence predictable) across a sequence of trials. They performed these search tasks both when the distractor location was not, and when it was predictable in advance. This allowed us to examine how distractor processing is shaped by expectations derived from spatial regularities across trials, both with and without expectations at the feature level. We found that both feature and spatial distractor regularities reduced distractor interference and affected post-distractor processing, as reflected in the amplitude and latency of the distractor-evoked Pd. No changes in pre-stimulus alpha-band activity were observed as a function of distractor location predictability. However, a control analysis revealed that counter to many studies where effects of distractor location probability on reaction time could not simply be explained by intertrial repetition (e.g., (Failing, Feldmann-Wustefeld, et al., 2019; Ferrante et al., 2018; Sauter et al., 2018), here the observed reduction in interference by distractors at the high probability distractor location largely reflected intertrial distractor repetition. This raises the possibility that some of our neural findings reflect intertrial priming, not spatial expectations. Therefore, before discussing our findings, we first discuss the natural conflation of intertrial priming and statistical learning (i.e., both contribute to performance in the same direction) and examine why effects of intertrial repetition were more pronounced here than in other studies, with at first sight, similar paradigms.

As the introduction of a high probability distractor location inevitably brings with it an imbalance in the probability of intertrial transitions, it is important to establish to what extent the observed suppression goes beyond intertrial distractor priming. To parse out the effect of intertrial transitions from the learning attributed to statistical regularities, a typical control is to repeat the analysis after eliminating all trials where the distractor location repeated, as we also did here. Although this allows one to parse out the independent contribution of intertrial priming, previous work suggests that effects of intertrial repetition reflect more than one mechanism (Feldmann-Wüstefeld & Schubö, 2016), and can become more pronounced in contexts where such repetitions are expected (van Moorselaar & Slagter, 2019; Won et al., 2019). Consistent with the notion that statistical learning can further increase the effect of intertrial priming, results of our exploratory linear mixed model analysis, in which multiple variables can be included in a single model and that thus allows for a more refined confound control, showed that the observed suppression in spatial bias blocks at the high probability distractor location could not solely be explained in terms of intertrial repetition. This finding corroborates previous studies showing that intertrial repetition effects in visual search are not solely stimulus driven, but can also reflect probabilistic expectations (Feldmann-Wüstefeld & Schubö, 2016; van Moorselaar & Slagter, 2019; Won et al., 2019).

Nevertheless, this still leaves unclear why the effect of statistical learning was much less pronounced here than in other studies and why this effect was not accompanied by impaired target processing at the high probability distractor location, as some studies have also reported (e.g., (Wang & Theeuwes, 2018b; Wang et al., 2019). One explanation may be found in the fact that counter to previous experiments, the high probability location was only introduced halfway through the experiment. It is therefore possible that by that time observers were less vigilant or more tired and thus less susceptible to any regularities across displays. Although we cannot exclude this scenario, we believe it unlikely as overall reaction times not only continued to decrease in the first, but also in the second half of the experiment, demonstrating that observer’s performance continued to improve. Alternatively, prior experience with unbiased situations may interfere with later statistical learning because observers are less prone to expect or use newly introduced probabilities once they already learned to efficiently deal with distractors, or because just as learned spatial biases persist after that bias is removed (Britton & Anderson, 2020; Ferrante et al., 2018; Sauter, Liesefeld, & Müller, 2019), the learned unbiased distribution also persists. In this respect it is interesting that in a pilot study (N = 24) with the same task and stimuli, where we directly introduced the high probability distractor location, we did find robust statistical learning using the standard intertrial subtraction control (see Appendix A). Future work needs to establish whether, and if so to what extent, statistical learning is indeed sensitive to prior exposure to unbiased conditions. Irrespective, for the remainder of this discussion it is important to consider that while our findings cannot be unequivocally contributed to either statistical learning or intertrial priming, the observed effects at high probability distractor locations to a large extent reflected intertrial repetitions.

We found that the ability to suppress distracting information was dependent on both distractor feature and location regularities, although large individual differences were observed in the extent to which individuals were sensitive to statistical regularities across trials. Extending previous studies (Burra & Kerzel, 2013; Gaspelin & Luck, 2018a), in no-spatial bias blocks, distractor interference was greatly reduced and associated with an early Pd and an earlier late Pd when distractor and target features were fixed and could thus be predicted compared to when they varied across trials. These findings suggest that distractors could be inhibited faster and more efficiently when distractor features were predictable in advance. Exploratory analyses suggested that in no-spatial bias blocks, these effects were dependent on developing feature-based expectations, as distractor costs were present at the start of each block and then gradually decreased, an effect that was accompanied by the emergence of the distractor-evoked Pd. In fixed-features blocks with spatial regularities, a Pd was evident already at the start of the block, indicating that spatial and feature expectations may interactively shape distractor suppression. There is some disagreement on the presence of distractor Pd in mixed-feature conditions, with some studies reporting an N2pc confirming attentional capture by the singleton distractors (Burra & Kerzel, 2013; Hickey, McDonald, & Theeuwes, 2006), whereas other studies observed a Pd albeit selectively on fast RT trials (McDonald et al., 2013). Here, at least on a group level, we observed a positivity that peaked relatively late, suggestive of a delayed late Pd in mixed-feature blocks without spatial regularities. As in the Wang et al. (2019) study, however, an early and late Pd emerged at the high probability distractor location, suggesting that suppressive mechanisms may onset earlier in time when expectations can be formed at the spatial level. Moreover, the early and late distractor-evoked Pd no longer differed in amplitude or latency between the mixed- and fixed-features conditions.

As discussed above, however, at the group level, the decrease in distractor interference in spatial bias blocks observed at the behavioral level was largely driven by intertrial repetitions, raising the possibility that the observed Pd effects (in part) reflect intertrial distractor location priming. In many studies reporting a distractor Pd, the Pd can be separated into an early and a late component (e.g., (Wang et al., 2019; Weaver et al., 2017). Whereas initial reports linked the early Pd (also referred to as PPC or lateralized P1) to lateral asymmetries in stimulus processing (Luck & Hillyard, 1994a) or saliency driven selection (Jannati et al., 2013), later studies suggest that the early Pd contributes to active suppression (Gokce et al., 2014), possibly via intertrial priming (Gokce et al., 2014). It is notable in this regard that EEG studies on statistical learning (e.g., (Wang et al., 2019), as was done here, only control for intertrial priming in the behavioral analysis, but not in the EEG analysis, leaving it unclear to what extent the observed EEG dynamics reflect pure statistical learning. That is, even when expectation-based suppression goes beyond intertrial distractor priming, this does not mean that the contribution of intertrial priming is negligible. Our exploratory analysis in which we distinguish between ‘learners’ (i.e., participants with impaired target processing at the high probability distractor location, and distractor filtering benefits above and beyond intertrial repetitions) and ‘non-learners’ (i.e., participants where target processing was not slowed down at the high probability distractor location, and intertrial repetitions could account for the observed distractor suppression benefits) may provide some insight into this question. Post-hoc analyses of the ERP data in spatial bias blocks showed that ‘non-learners’ only exhibited the early distractor Pd, while, although not reaching statistical significance, ‘learners’ only exhibited the typical late Pd. Albeit speculative, given the post-hoc nature of these latter findings and the small samples sizes of the two groups, the early Pd may hence reflect changes in sensory processing related to intertrial priming, possibly in retinotopically organized visual regions (given that the eyes were at fixation at all times), while the late Pd may reflect suppression related to spatial statistical learning at a higher level in the processing hierarchy, where information is integrated over a longer time scale, at the spatiotopic level. It is notable in this regard that clinical studies in neglect patients also suggest that intertrial priming through distractor repetition operates earlier in the visual processing hierarchy (Finke et al., 2009; Saevarsson, Jóelsdóttir, Hjaltason, & Kristjánsson, 2008). The early Pd may by itself be insufficient to reduce distractor interference as in subjects that did not show impaired target processing at high probability distractor locations (‘non-learners’), we only observed an early Pd to targets. Future research is necessary to more conclusively determine the functional significance of the early and late Pd, and to more systematically examine individual differences in statistical learning.

While it is typically assumed that pre-attentive suppression as indexed by the (early) Pd relies upon proactive inhibition, to date very few studies have also examined anticipatory markers of suppression. Of the few studies that did, despite the prevailing view that alpha oscillations implement active top-down inhibition (Jensen & Mazaheri, 2010), only a single study observed an increase in anticipatory alpha-band activity contralateral to the likely distractor location that was followed by a subsequent Pd to any stimulus presented at that location (Wang et al., 2019). By contrast, two other studies observed expectation-based distractor suppression as indexed via (early) Pd modulations in the absence of any distractor location-related anticipatory alpha lateralization (Noonan et al., 2016; van Moorselaar & Slagter, 2019). A notable difference that might explain this discrepancy in findings is that in these latter two studies, distractor (and target) features were fixed, whereas in the former study they varied from trial to trial, rendering it possible that generic space-based proactive suppression only develops when distractors cannot be predicted at the feature level. While the current study was designed to address this possibility, the present results do not allow for drawing any resolute conclusions as counter to Wang et al. (2019), in the spatial bias blocks the observed distractor benefits only marginally went above and beyond intertrial priming, nor were they accompanied by impaired target processing. These findings are not indicative of generic space-based proactive suppression, leaving it unclear whether anticipatory alpha-band modulations implement generic space-based proactive suppression or not. Yet, even the participants that were most sensitive to spatial statistical learning did not show any modulation of anticipatory alpha activity. Although one should be careful in interpreting null findings and there may be alternative explanations why alpha modulations were absent in ‘learners’, the current findings add to a rapidly growing literature suggesting that alpha-band oscillations do not appear to play a causal role in proactive distractor suppression (Antonov, Chakravarthi, & Andersen, 2020; Foster & Awh, 2018).

A notable aspect of the current findings is that spatial and feature expectations were associated with a larger early Pd and/or an earlier-onset late Pd. This pattern appears to be at odds with an influential explanation of expectation-based distractor suppression, predictive coding (Friston, 2009; Noonan et al., 2018) which posits that previous experiences with distracting information attenuates sensory processing. Within the predictive processing framework, current sensory input is compared to stored representations that are grounded in the expected frequency and contextual properties of distractors, and the greater the match between the two, the more the sensory response is attenuated or explained away. In this scenario, one would thus predict repeated distractors to evoke smaller responses and elicit less interference in the system, reducing the need for distractor inhibition. Consistent with this, several previous studies have reported that the amplitude of the Pd decreases as a function of expectations (Heuer & Schubö, 2019; van Moorselaar, Lampers, Cordesius, & Slagter, 2020; van Moorselaar & Slagter, 2019). At face value such a reduced Pd also appears consistent with habituation (Thompson, 2009; Turatto & Pascucci, 2016), the progressive attenuation of a response to repeated sensory stimulation, which is intimately linked to predictive coding (Chelazzi, Marini, Pascucci, & Turatto, 2019). However, habituation specifically refers to the decrease in strength of an orienting response and it could thus also be argued that an increased and/or earlier Pd reflects a more efficient re-orienting following initial attentional capture by the distractor (Moher & Egeth, 2012). While such a perspective is in line with the present pattern of results, it is for example at odds with the pattern reported in the Wang et al. (2019) study where the observed Pd effect was accompanied by proactive suppression as indexed by impaired target processing at the suppressed location and anticipatory alpha modulations contralateral to the high probability distractor location. To date, within the current theoretical frameworks it thus remains unclear why in certain contexts, as in the present study, the Pd increases, whereas in others, it decreases and future work is necessary to explain this apparent discrepancy.

To conclude, here we find that the ability to inhibit distracting information is strongly affected by both feature-based and space-based distractor regularities, as indicated by corresponding reductions in distractor costs during visual search and in the amplitude and latency of the distractor-evoked Pd ERP component. Our findings also raise the intriguing possibility that current distractor learning may be influenced by the preceding task context, as we found that while in spatial bias blocks preceded by no-spatial bias blocks, observers were sensitive to regularities across longer time scales to some extent, the observed effects of distractor location probability largely reflected intertrial repetition. Suggestive of effects of both intertrial priming and probabilistic expectations on distractor processing, exploratory analyses revealed individual differences in sensitivity to distractor location regularities that may suggest a functional dissociation between the early and late Pd: while the early Pd may reflect early sensory suppression related to intertrial priming, the late Pd may reflect later spatiotopic suppression related to statistical learning over a longer time scale. Together these findings reveal how previous encounters with distracting information shape distractor processing at both the feature and the spatial level.

## Acknowledgements

This research was supported by a European Research Council (ERC) starting grant (679399) to H.A.S. D.v.M. contributed to design, collected the data, performed the analyses, and contributed most of the writing. HS was closely involved in the design of the experiment and the analysis plan and made significant contributions to the writing. ND collected the data and provided feedback on the writing. We would like to thank Mingwei Zhou for her valuable help with data collection. We would like to thank dr. Heinrich R. Liesefeld for his valuable and constructive comments throughout the review process.

## Appendix A. Supplementary data

Prior to the EEG experiment, we conducted a behavioral pilot experiment specifically aimed to establish whether we could replicate two key characteristics of statistical learning as reported in the literature: 1) robust suppression at high probability distractor locations after eliminating from the data all intertrial transitions, and 2) impaired target processing at high probability distractor locations in mixed-feature, but not in fixed-feature conditions. Yet, in contrast to the EEG experiment, the spatial bias was introduced already at the beginning of the block. As the findings of this pilot study are of interest for our speculation why these markers were less robust (1) and absent (2) in our subsequent EEG study, where the spatial regularities were only introduced half-way into the experiment, we include them here in support of our speculation that prior experience with neutral blocks may interfere with later statistical learning.

### Methods

The methods and analyses (https://osf.io/53pnk/) were the same as in the main EEG experiment, except for the following changes. A planned number of 24 participants (Mean age = 21 years, range 19 – 26; eleven men) participated in the experiment. Four participants were replaced because either their error rate and/or average RT were more than 2.5 SD from the group mean. Each trial started with a 750-ms black fixation display containing only a white rim. Also, counter to the main experiment, targets and distractor locations were not limited to the vertical and horizontal axis, but instead could appear on all eight locations in the search display. Targets appeared with equal probability on all locations, whereas distractors, when present (68% of trials), appeared with a higher probability (63%) at one of the eight locations (counterbalanced across participants).

Critically, within a single experimental session, the dynamics of consecutive search displays were changed in four separate conditions. In addition to mixed- and fixed-feature conditions, identical to the main experiment, we included shape-fixed and color-fixed condition blocks (order counterbalanced across participants), in which, respectively, the shape configuration was fixed with random varying colors or vice versa.

Participants completed six blocks of 56 trials each for each condition (order counterbalanced). Each sequence of six blocks was preceded with 56 practice trials without a high probability distractor location manipulation. At the start of each block participants received written instructions about the dynamics of the upcoming set of trials (e.g., ‘In the upcoming block the target and distractor will have random shapes and colors).

### Results and Discussion

Exclusion of incorrect responses (4.9%) and data trimming (2.9%) resulted in an overall loss of 7.8% of the data. As visualized in Figure 1, and consistent with the main experiment, a repeated measures ANOVA with within subjects’ factors Condition (predictable, shape predictable, color predictable, unpredictable) and Distractor location (high probability distractor location, low probability distractor location) yielded a main effect of Distractor location (*F* (1, 23) = 57.4, *p* < 0.001, 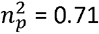) reflecting reduced interference at high probability distractor locations. This main effect was accompanied by an interaction showing that the benefit at the high probability distractor locations was most pronounced in mixed-feature and color-fixed conditions versus fixed-feature and shape-fixed conditions (*F* (3, 69) = 2.9, *p* = 0.04, 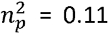). Nevertheless, distractors continued to produce reliable interference at both the low (all *t*’s > 6.5, all *p*’s < 0.001, all *d*’s > 1.3) and high (all *t*’s > 2.3, all *p*’s < 0.029, all *d*’s > 0.48) probability locations across all conditions. Counter to a suppressive gradient centered on the high probability distractor location as reported elsewhere (e.g., (Wang & Theeuwes, 2018b), however, an analysis with Condition and Distance from high probability location (ranging from 1 to 4; thus, excluding the high probability location itself) showed no main effect of Distance nor an interaction (all *F*’s < 1.4, all *p*’s > 0.25).

**Figure 1.**
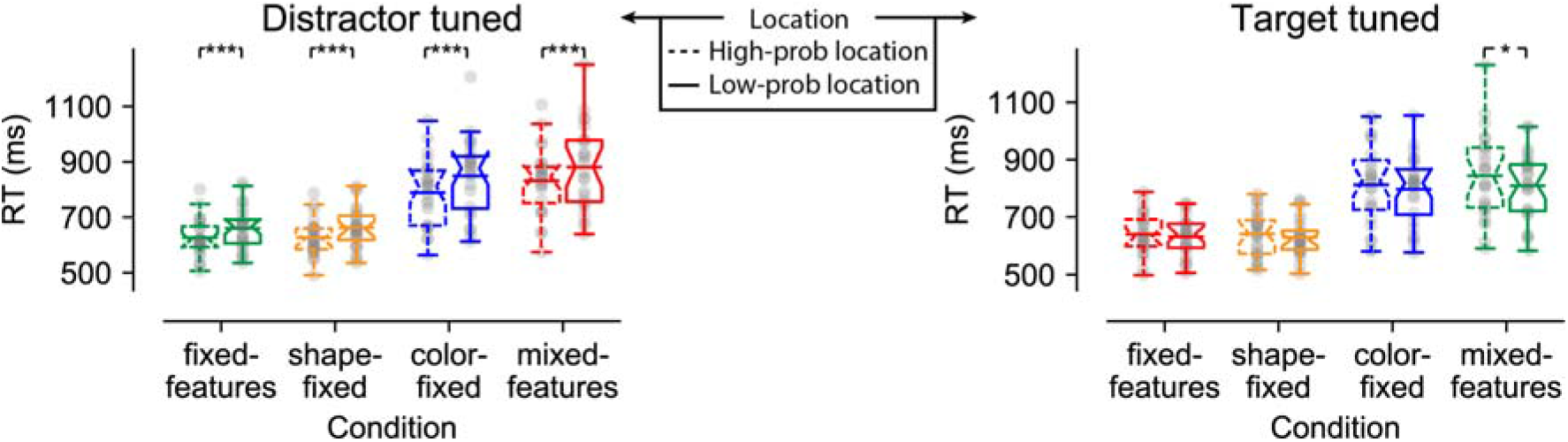
Behavioral findings visualized by notched boxplots with solid horizontal lines corresponding to the mean. (A) Reaction times as a function of distractor location (left) and target location (right) across conditions.

Critically, and counter to the main experiment, the main effect of Distractor location (*F* (1, 23) = 47.2, *p* < 0.001, 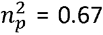) remained reliable after removal of all trials where the distractor location repeated from the data (i.e., standard intertrial priming control). Across all conditions RTs were faster for distractors at high probability distractor locations than at low probability distractor locations (all *t*’s > 4.4, all *p*’s < 0.001, all *d*’s > 0.90). At the same time the observed interaction was no longer significant (*F* (2, 43) = 1.8, *p* = 0.15, 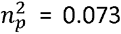), suggesting that larger suppressive benefits observed in the mixed-feature and color-fixed conditions to a large extent could be explained by stronger intertrial priming effects in these conditions.

After having established robust distractor suppression at the high probability distractor location, we examined whether target processing was also impaired at this location. As in distractor absent trials the number of observations at high probability locations was relatively low (N = ~20), to increase power we also included distractor present trials in this analysis, but, to prevent a confound in the analysis, only those trials where the distractor appeared at a low probability location. Consistent with reports of feature blind suppression at high probability distractor locations, this analysis yielded a main effect of Target location reflecting slower RTs at high probability distractor locations (*F* (1, 23) = 6.4, *p* = 0.019, 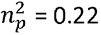), and this effect remained reliable after exclusion of all trials where either the distractor or the target location was primed (*F* (1, 23) = 4.7, *p* = 0.041, 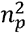 = 0.17). Also, consistent with the idea that generic spatial suppression only develops when the spatial distractor regularity cannot be combined with feature expectations (Allenmark et al., 2019; Sauter et al., 2018; Zhang et al., 2019), planned pairwise comparisons identified impaired target processing in the mixed-feature condition (*t* (23) = 2.9, *p* = 0.009, *d* = 0.58), the one condition without feature regularities, but not in the other conditions (all *t*’s < 1.4, all *p*’s > 0.17, all *d*’s < 0.29). Note however, that this dissociation was not confirmed by a significant interaction (*F* (2, 51) = 1.8, *p* = 0.18, 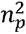 = 0.072).

Finally, we tested whether participants could correctly identify the high probability distractor location. Consistent with the idea that the observed suppressive effects, at least to a large extent, rely on implicit learning, only one out of 24 participants correctly identified the high probability distractor location (chance level = 12.5% or 3 participants) and the average deviation from the high probability location was 2.2 (chance level = 2).

These results confirm that, when a high probability distractor location is immediately introduced, distractors at this location are not only more efficiently ignored, but this effect clearly goes beyond intertrial distractor location priming. Second, the results appear to be in line with the observation that this spatial suppression is feature blind when learning can by design only occur at the feature level.

## Appendix B. Eye movement control analysis

**Figure 2.**
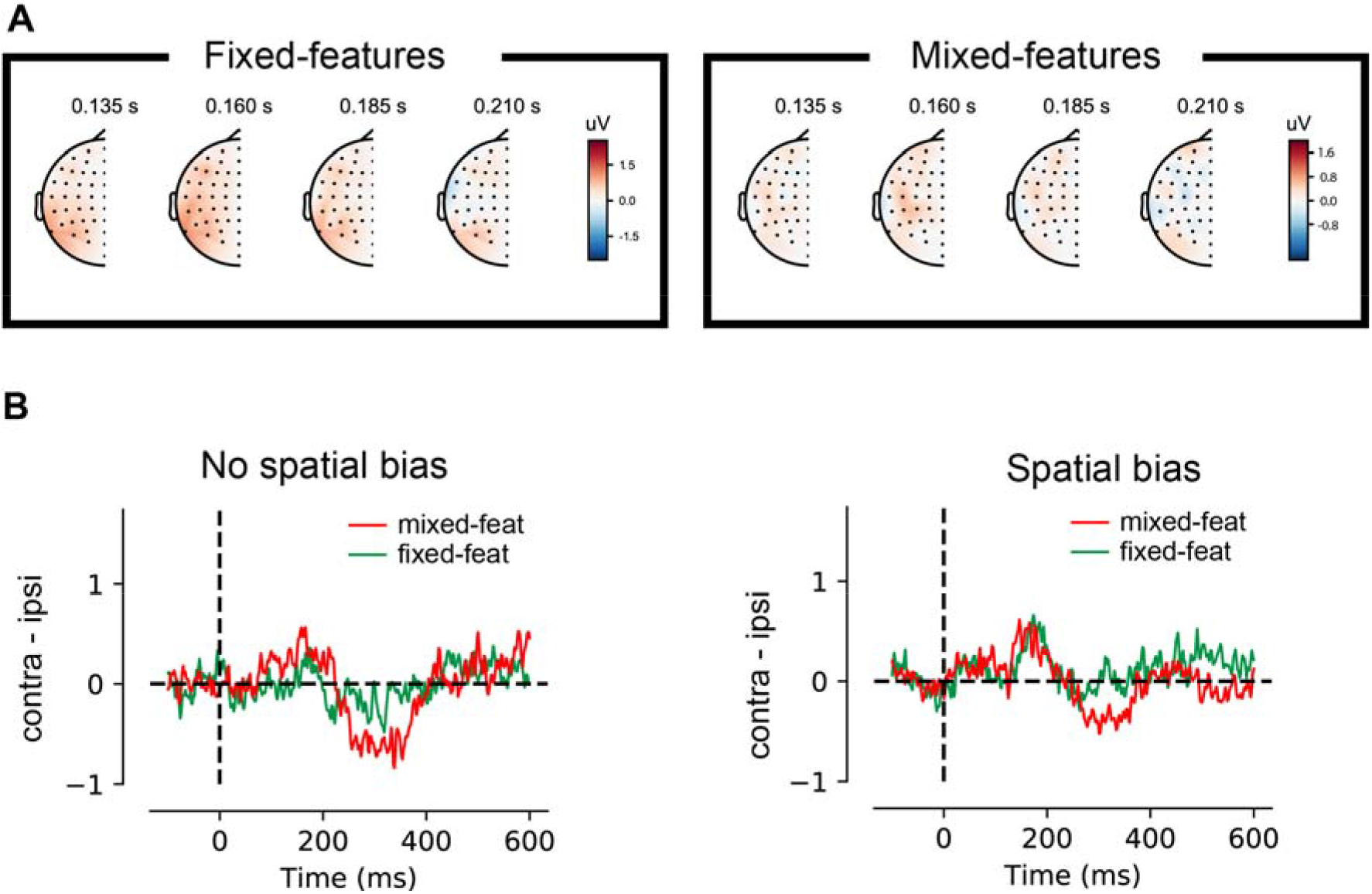
Eye movement control analysis. A) Topographic distribution for contra vs ipsilateral activity in no spatial bias blocks tuned to the distractor location. B) Contralateral vs ipsilateral hEOG traces relative to the distractor location across conditions in no spatial bias and spatial bias blocks. Although numerically there appeared to be a difference between mixed- and fixed-feature conditions, cluster-based permutation tests failed to identify conditions differences, nor did any of the traces reliably differ from zero. Previous research has shown that residual EOG activity of 3 V corresponds to an average eye movement of less than 0.2 degrees of visual angle and a propagated voltage of less than 0.1 V at the posterior scalp sites (Lins, Picton, Berg,& Scherg, 1993). Together these findings show that our eye-movement exclusion procedure was successful and the observed lateralized effects at posterior electrodes cannot be explained by systematic eye-movements.

## Footnotes

1 Due to a programming error, in five subjects the high probability location was introduced after 7 blocks and in another seven subjects after 9 blocks, instead of after eight blocks.

2 The distractor location beta weight remained negative (17.0) and significant (*p* = 0.005) when target position, which was counterbalanced over distractor present and absent trials (Failing, Wang, & Theeuwes, 2019; Zhang et al., 2019), was included as a control variable in the model.

3 Note, that the same subjects would have been selected in case we used the corrected target-location effect (i.e., after controlling for four different forms of intertrial priming (D_n-1_ – D_n_, D_n-1_ – T_n_, T_n-1_ – D_n_, T_n-1_ – T_n_)).

4 Here we chose peak-latency rather than area latency to minimize the risk that the analysis was confounded by overlapping activity from the preceding early Pd component (Heinrich René Liesefeld, 2018).

## Notes

Conflict of Interest: The authors declare no competing financial interests

### Competing Interest Statement

The authors have declared no competing interest.

### Summary of Updates

Minor updates after fourth round of review

